# Human mitochondrial transfer modeling reveals biased delivery from mesenchymal-to-hematopoietic stem cells

**DOI:** 10.1101/2025.01.06.631461

**Authors:** Steven J. Dupard, Joao M. Pinheiro, Aurélie Baudet, Alejandro Garcia Garcia, Dimitra Zacharaki, David Hidalgo Gil, Sujeethkumar Prithiviraj, Vinay Swaminathan, Paul E. Bourgine

## Abstract

Within the bone marrow (BM), the intercellular communication between hematopoietic stem and progenitor cells (HSPCs) and mesenchymal stem/stromal cells (MSCs) is critical for the life-long maintenance of functional hematopoiesis. In recent years, the transfer of mitochondria between MSCs and HSPCs has emerged as a key aspect of this communication, occurring both in stress and homeostatic conditions. In human, the mesenchymal-to-hematopoietic transfer process and functional impact remain cryptic, primarily due to a lack of robust models. To this end, we here describe the development and exploitation of iMSOD-mito, an immortalized human MSCs line bearing an inducible mCherry mitochondrial tag. Co-culture with primary healthy HSPCs or a leukemic cell line revealed a high mitochondrial transfer rate (>15%), exclusively relying on cell-to-cell contact. While all CD34+ blood cells received mitochondria, a preferential transfer towards phenotypic hematopoietic stem cells was identified. Similarly, using primary MSCs with genetically labelled mitochondria we confirmed a transfer to all CD34+ populations, albeit occurring at a lower frequency than with the iMSOD-mito (3.38%). By engineering 3D bone marrow niches in perfusion bioreactor, this transfer rate could be significantly increased, while the biased towards HSC as receiver was maintained. Functionally, mitochondria-receiving cells exhibited an increased mitochondria membrane potential and reactive oxygen species (ROS) production, which in HSPCs was associated with retained quiescence in single cell divisional assay. In summary, we propose the iMSOD-mito as a standardized tool to model human mesenchymal-to-hematopoietic mitochondria transfer in 2D or 3D culture systems. Our work prompts the study of mitochondria transfer in both healthy or disease conditions, towards the design of regenerative therapies or identification of new targets in a malignant context.

## 1 Introduction

Cell-to-cell mitochondrial transfer, also referred to as horizontal mitochondrial transfer, was first evidenced in vitro by the restoration of mitochondrial activity of mitochondria deficient A549 *p°* cancer cells (Spees *et al*., 2006) and in vivo by the mitochondrial DNA (mtDNA) polymorphism observed in transmissible canine venereal tumor (Rebbeck, Leroi and Burt, 2011). Since these pioneering reports, such transfer was observed in a diversity of tissue as a rescue mechanism in pathological conditions affecting mitochondrial functions (Islam *et al*., 2012; Vallabhaneni, Haller and Dumler, 2012; Hayakawa *et al*., 2016), to the benefit of involved cells. More importantly, engineered inter-mitochondrial mouse chimeras, in which tissue is formed from two cell lineages with divergent mtDNA haplotype, revealed mitochondrial transfer as a fundamental physiological process during normal development and maintenance of all adult tissues (Marti Gutierrez *et al*., 2022).

Bone marrow mesenchymal stem/stromal cells (BM-MSCs) deliver mitochondria to neighboring cells through this process. BM-MSCs are BM niche resident cells responsible for the maintenance and regulation of hematopoiesis via secreted factors and direct cell-cell interactions with hematopoietic stem and progenitor cells (HSPCs) (Mendelson and Frenette, 2014; Wei and Frenette, 2018). As niche entity, BM-MSCs possess extensive regenerative and anti-inflammatory functions which makes them a popular cell source for cell-based therapies (Wei *et al*., 2013). As such, their exploitation in disease models exposed their mitochondrial transfer capabilities, the impact of which has been extensively studied across a variety of tissues. In mice, BM-MSCs mitochondrial transfer was shown to be essential to the immunomodulatory and anti-inflammatory role of recipient cells. Among others, BM-MSCs in vivo mitochondria transfer alleviates macrophage inflammatory phenotype in diabetic nephropathy (Yuan *et al*., 2021), restore proliferation and metabolic activity in alveolar epithelial cells during lung infection (Islam *et al*., 2012), and increase the survival of motor neuron in a model of spinal cord injury (Li *et al*., 2019). Not surprisingly, BM-MSCs mitochondria transfer was also observed in BM, where transfer to HSPCs is involved for the survival from bacterial infection or myeloablation (Mistry *et al*., 2019; Golan *et al*., 2020). In these conditions, disruption of the transfer between BM-MSCs and HSPCs results in a steep increase in animal mortality, exemplifying the importance of this communication route for hematopoietic functions.

Naturally, human BM-MSCs cells were also investigated for their capacity to transfer mitochondria. Much like their murine counterparts, human BM-MSCs transfer was associated with improved metabolic functions of injured cells in mouse xenograft models (Islam *et al*., 2012; Tseng *et al*., 2021). Importantly, human BM-MSCs mitochondrial transfer plays a pivotal role in drug-resistance in leukemia, with in vitro evidence for acute myeloid leukemia (Wei *et al*., 2013; Moschoi *et al*., 2016; Mistry *et al*., 2021; Saito *et al*., 2021; Cai *et al*., 2022), T-cell acute lymphoblastic leukemia (Wang *et al*., 2018), B-cell precursor acute lymphoblastic leukemia (Wang *et al*., 2018) and multiple myeloma (Marlein *et al*., 2019; Giallongo *et al*., 2022).

While evidence is still scarce, one can presume that mitochondrial transfer from BM-MSCs also contributes to normal hematopoiesis. Previous reports demonstrated mitochondria transfer from BM-MSCs to CD34+ cells (Moschoi *et al*., 2016; Marlein *et al*., 2017) and identified that subpopulations of CD34+, namely CD3+ and CD3-, do not receive mitochondria to the same extent (Moschoi *et al*., 2016). This shows not only that non-malignant hematopoietic cells receive mitochondria from BM-MSCs but also suggests that a selective mechanism underlies the targeting of hematopoietic populations to be transferred, in correlation with the phenotype of recipient cells. However, despite evidence for transfer to normal hematopoietic cells, this research axis was not further explored.

While correlative, multiple insights contributed to our understanding of the mechanism of mitochondrial transfer by BM-MSCs. Notably, the role of gap junctions facilitating this transfer in conjunction to AKT-PI3K pathways activation and the CD38 expression on recipient cells (Islam *et al*., 2012; Marlein *et al*., 2019; Mistry *et al*., 2019). However, multiple caveats are still limiting the exploitation of these studies. Indeed, findings are bound to a particular tissue or disease model and does not reflect the ubiquitous nature of mitochondrial transfer in tissue maintenance (Marti Gutierrez *et al*., 2022). Furthermore, inhibition of the molecular players identified does not completely abrogate mitochondrial transfer, pointing at a level of redundancy in pathways leading to mitochondrial transfer yet to be uncovered. Importantly, the number of human-only mitochondrial transfer reports are scarce which limits translational applications of this mechanism. A possible culprit might relate to the extensive use of Mitotracker and similar dyes for tracking human mitochondrial transfer disrupting mitochondria and excreted by xenobiotic pumps. These dyes were indeed recently identified to be inconsistent in labelling normal HSPCs (de Almeida *et al*., 2017; Mansell *et al*., 2021). As the significance of human mitochondrial transfer unravels, a human-only standardized protocol is needed to validate key molecular and cellular players for exploitation, which the model presented hereafter aims at providing.

Here, we aim at generating and exploiting an immortalized human BM-MSC for a dye-independent tracking of mitochondrial transfer towards malignant and normal human HSPCs. This exploitation aims at identifying discriminants and consequences of transfer to normal human blood cells in a standardized manner.

## Results

### iMSOD-mito is a human mesenchymal cell line with robust mitochondrial transfer capacity to human hematopoietic cells

The standardization of mesenchymal-to-hematopoietic mitochondrial transfer would largely benefit from the use of a dedicated cell line. Using the murine mesenchymal line OP9, we first observed an effective transfer of mitochondria following 48h of co-culture with human umbilical cord blood (UCB) CD34+ hematopoietic (**Supp. Figure 1A**). This was detected by species-specific mitochondrial DNA analysis, thus confirming effective mouse to human transfer.

To study the process in a human specific setting, we targeted the engineering of of a pre-existing BM-MSC line defined as Mesenchymal Sword of Damocles (MSOD). The MSOD cells were transduced with a genetic cassette consisting in a doxycycline-inducible (TetO) mitochondrial localization signal tagged to mCherry, coupled with the trCD8a surface marker (**Figure 1A**). Following transduction, the resulting iMSOD-mito exhibited clear mCherry mitochondrial labelling upon doxycycline induction (**Figure 1B**), confirmed by colocalization with the mitochondrial import receptor subunit TOMM20 using confocal microscopy (**Supp. Figure 1B**). The iMSOD-mito also displayed uniform expression of trCD8 (98% ± 1.21, **Supp. Figure 1C**) when exposed to doxycycline, validating the full functionality of the implemented genetic device.

**Figure 1.**
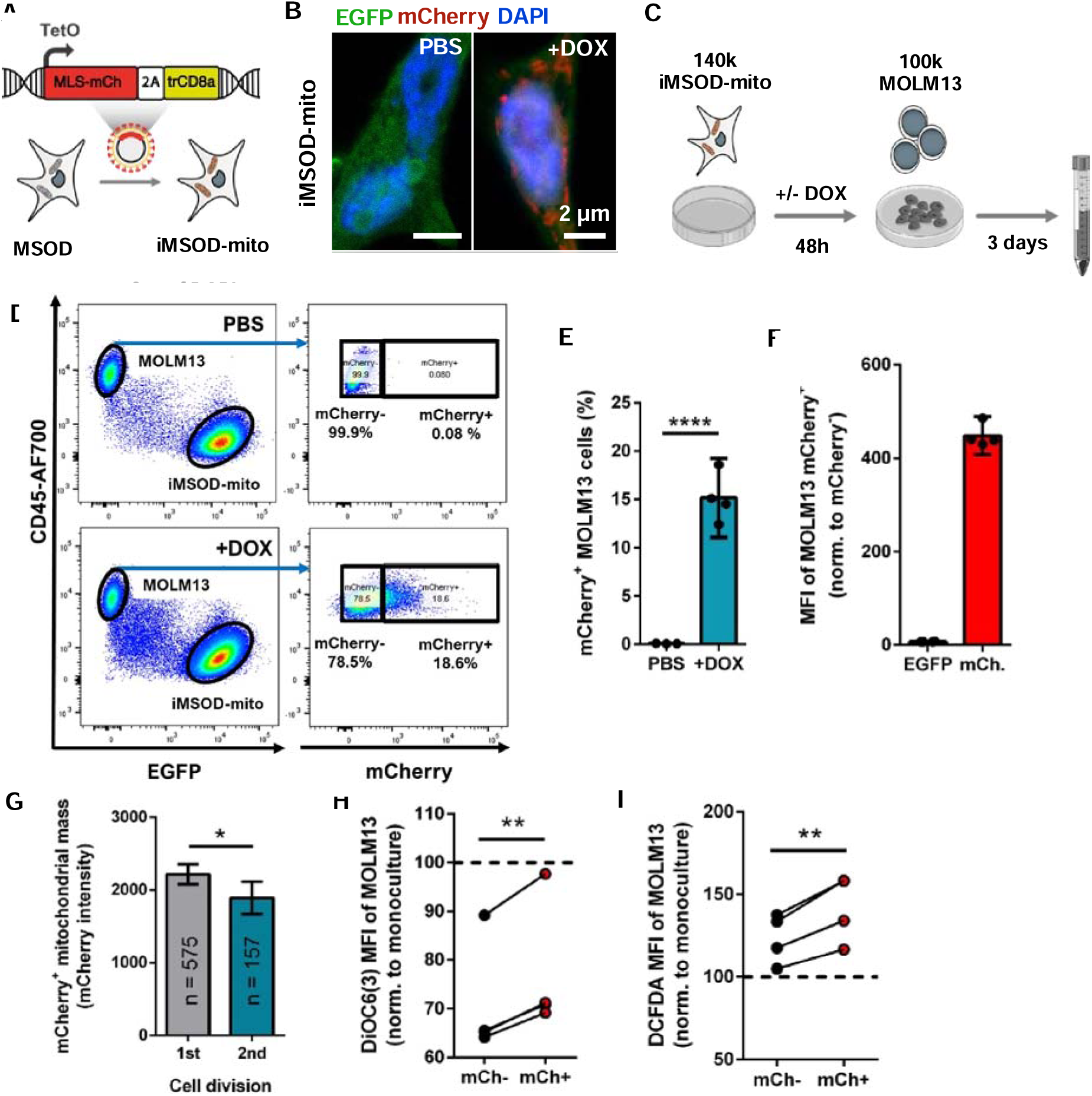
iMSOD-mito is a human mesenchymal cell line with robust mitochondrial transfer capacity to human hematopoietic cells. **(A)** Diagram representing the genetic cassette transduced to the human bone marrow mesenchymal sword of Damocles (MSOD) cell for the generation of the inducible mitochondria mCherry (mCh) tagged iMSOD-mito. MSL: mitochondrial localization signal. **(B)** Confocal microscopy of iMSOD-mito after 24h exposure to phosphate-buffered saline (PBS) or 150 ng/mL of doxycycline (DOX). **(C)** Experimental scheme of the coculture of MOLM13 cells on a confluent layer of iMSOD-mito with or without induction of mitochondrial mCherry tagging. **(D)** Flow cytometry gating of MOLM13 and iMSOD analysis of the mCherry intensity in MOLM13 cells after three days of coculture**. (E)** Percentage of mCherry+ MOLM13 cells after coculture with (+DOX) or without (PBS) induction of iMSOD-mito. Unpaired t-test with logit transformation (n = 4). **(F)** Mean fluorescence intensities (MFIs) of mCherry and EGFP of mCherry+ MOLM13 cells, normalized to mCherry-MOLM13 MFIs. (n = 4). **(G)** Mitochondrial mass of mCherry+ MOLM13 according to their divisional history after coculture with iMSOD-mito, determined with CellTrace Violet (gating in Supp. Figure 1D). Welch’s t-test (n_1_= 575, n_2_= 157). **(H)** Mitochondrial membrane potential of mCherry+ and mCherry-MOLM13 measured by DiOC6(3) MFI after coculture with iMSOD-mito. Data are normalized to MOLM13 monoculture DiOC6(3) MFI (dashed line). Paired t-test (n = 4). **(I)** Reactive oxygen species activity in mCherry+ and mCherry-MOLM13 measured by DCFDA MFI after coculture with iMSOD-mito. Data are normalized to MOLM13 monoculture DCFDA MFI (dashed line). Paired t-test (n = 4). **** = *p*-value <.0001. ** = *p*-value <.01. * = *p*-value <.05.

We next aimed at evaluating the mitochondria transfer capacity of the iMSOD-mito line towards human hematopoietic cells. To this end, MOLM13, an acute myeloid leukemia cell line previously reported to be a recipient of mitochondria from BM-MSCs (Saito *et al*., 2021), were co-cultured with iMSOD-mito for 3 days in presence of doxycycline (or PBS, control) (**Figure 1C**). Following cell retrieval, flow cytometry allowed the clear identification of the two populations with MOLM13 expressing the pan leukocyte CD45 marker and the iMSOD-mito being EGFP positive. (**Figure 1D**, **Supp. Figure 1B**). In presence of doxycycline, a significant fraction of the MOLM13 cells received mCherry mitochondria from iMSOD-mito (15.15% ± 2.57, **Figure 1E**). Of interest, within the mCherry+ MOLM13 population, the mean fluorescence intensity (MFI) of EGFP was unchanged compared to mCherry-MOLM13 suggesting that no-to-limited cytoplasmic elements were transferred alongside the mitochondria (**Figure 1F**).

By correlating MOLM13 cell divisional history and their mitochondrial mass (mCherry mean fluorescence intensity), we observed a slight but significant decrease of mCherry mitochondria in cells that underwent two divisions. This suggests a continuous rather than discrete mitochondria transfer also occurring in daughter cells (**Figure 1G, Supp. Figure 1D**). Mitochondrial membrane potential (MMP), measured by DiOC6(3), did not significantly change in MOLM13 in coculture with iMSOD-mito (**Supp. Figure 1E**). Moreover, the induction of mCherry labeling of mitochondria and the presence of DOX in the culture media did not affect the MMP of MOLM13 cells (**Supp. Figure 1E).** This indicates that neither the coculture setting nor the presence of doxycycline affects mitochondrial activity in MOLM13. However, mCherry+ MOLM13 displayed a greater MPP and reactive oxygen species (ROS) level than mCherry-MOLM13, pointing at a higher metabolic activity in recipient cells (**Figure 1H** and **1I**) and the transfer of functional mitochondria.

Lastly, to investigate the transfer between MSCs, uninduced iMSOD-mito stained with CellTrace Far Red (CTFR+) were placed in coculture for 24h with unstained (CTFR-) doxycycline-induced iMSOD-mito. The presence of mCherry in CTFR+ iMSOD-mito will thus report the rate of mitochondrial transfer. We observed transfer to 1.02% (± 0.13) of CTFR+ iMSOD-mito (**Supp. Figure 1F-G**), thus validating the presence of transfer between MSCs.

In summary, we here report the iMSOD-mito as human BM-MSCs cell line capable of transferring functional mitochondria to human hematopoietic cells.

### iMSOD-mito transfer mitochondria exclusively via cell-to-cell contact in a non-stochastic manner

Mitochondria transfer has been reported to potentially occur through cell-to-cell contact, extracellular vesicles, or free-floating mitochondria (Borcherding and Brestoff, 2023). To identify the mode of transfer from iMSOD-mito, we established various culture conditions with MOLM13 (**Supp. Figure S2A**). As anticipated, direct contact via standard co-culture condition led to significant mitochondria transfer (**Figure 2A**). In sharp contrast, co-culture in transwell or conditioned media collected from iMSOD-mito did not lead to a detectable transfer of mitochondria in MOLM13 (**Figure 2A**). This points at a cell-contact driven transfer of mitochondria, possibly relying on the formation of tunneling nanotubes (TNT). Using time-lapse microscopy, we confirmed the mesenchymal-to-hematopoietic transfer of mitochondria (**Figure 2B**). The captured process was initiated by close cell membrane contact, with the formation of short-range TNTs between iMSOD-mito and MOLM13 leading to mCherry mitochondria transfer to MOLM13 within a 40-minute timeframe.

**Figure 2.**
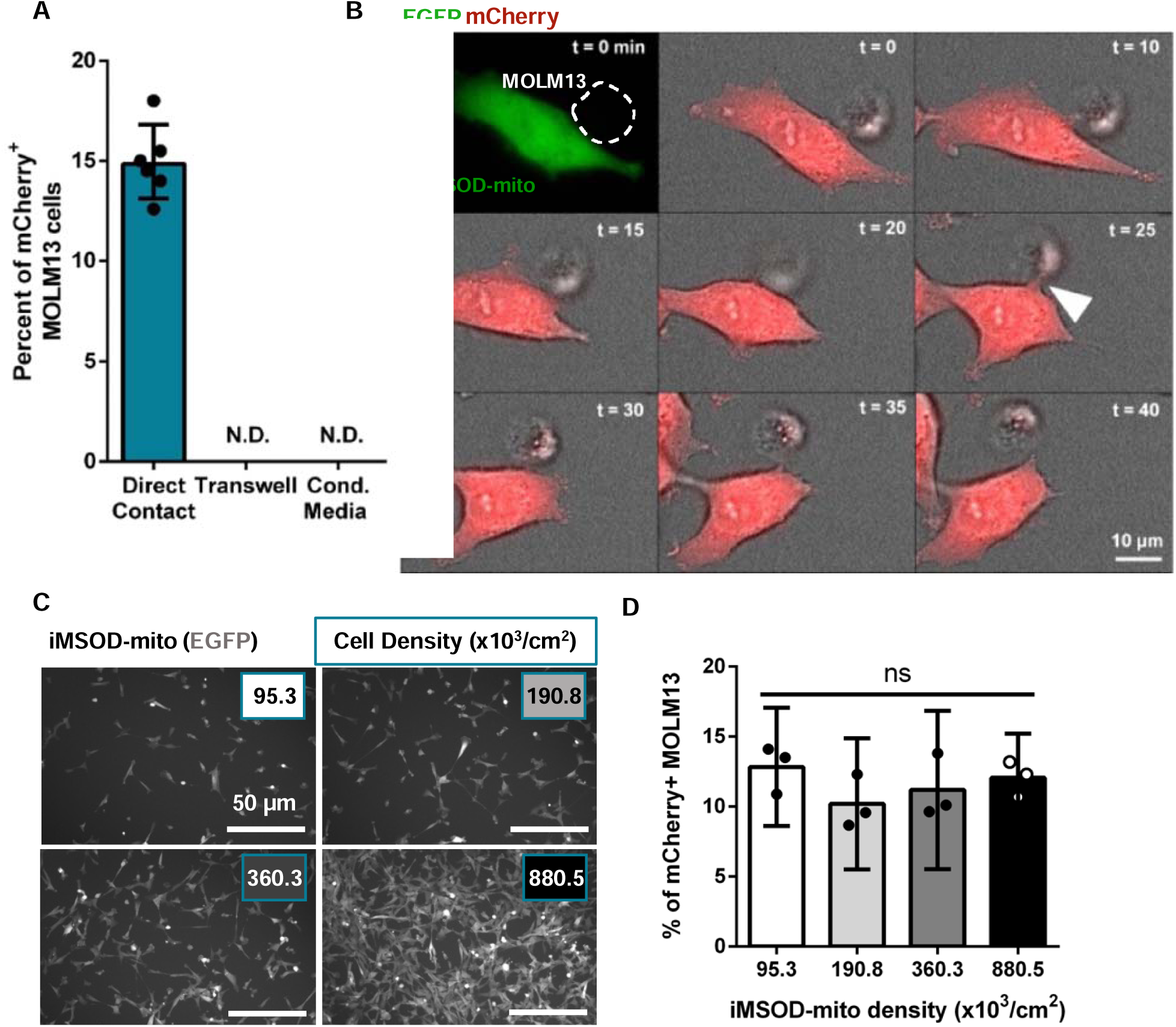
iMSOD-mito transfer mitochondria exclusively via cell-to-cell contact in a non-stochastic manner. **(A)** Percentage of mCherry+ MOLM13 cells after 3 days coculture with induced iMSOD-mito in direct contact, through a transwell or exposed to induced iMSOD-mito conditioned (cond.) media (n = 6). N.D.: Not detected. **(B)** Live imaging snapshots of the mitochondrial transfer from induced iMSOD-mito (EGFP and mCherry) to MOLM13 (dashed contour). After cell-cell contact is established, mCherry mitochondria transfer is initiated at t = 25 min (arrowhead). **(C)** Confocal microscopy analysis of induced iMSOD-mito density prior to MOLM13 coculture. iMSOD-mito density was determined from manual counting of three fields of view. **(D)** Percentage of mCherry+ MOLM13 cells after 3 days coculture with induced iMSOD-mito at different densities. One-way ANOVA (n = 3). ns = *p*-value >.05.

As this transfer involved direct cell-cell contact, we questioned whether the transfer efficacy could be influenced by the mesenchymal density in our culture system. To this end, MOLM13 were co-cultured with four different iMSOD-mito cell density, corresponding to a 2-fold, 4-fold and 8-fold increase respectively (**Figure 2C**). Importantly, the mesenchymal density difference was maintained throughout the culture time, as assessed by the constant iMSOD-mito doubling time across conditions (**Supp. Figure 2B**). Surprisingly, the percentage of MOLM13 being transferred remained unaffected by the iMSOD-mito culture densities (**Figure 2D**). This suggests that the mitochondrial transfer is not guided by stochastic interactions between MOLM13 and iMSOD-mito.

In brief, we here show that mitochondrial transfer from iMSOD-mito to MOLM13 exclusively relies on cell-to-cell interactions and establishment of short-range TNTs.

### iMSOD-mito transfer mitochondria preferentially to phenotypically stem human HSPCs

We next investigated the capacity of iMSOD-mito to transfer mitochondria to primary, healthy hematopoietic populations. UCB-CD34+ HSPCs were thus isolated and co-cultured with iMSOD-mito (**Figure 3A**). The presence of mCherry-labelled mitochondria was clearly detected in retrieved CD34+ cells (**Figure 3B** and **3C**), with over 8.5% (± 4.41) of cells receiving mitochondria. We first observed no difference in terms of transfer frequency within the pan hematopoietic marker CD45+ and the stem and progenitor marker CD34+ populations (**Supp. Figure 3A**). To identify a potential variable transfer within hematopoietic sub-populations, we assessed the mCherry+ frequency in stem and progenitor cells (**Supp. Figure 3B**). This revealed an unequal mitochondrial transfer across hematopoietic phenotypes (**Figure 3D**). The CD34+CD45ra-(6.81% ± 3.76), and both the CD34+CD45ra-CD90-EPCR+ (5.07% ± 4.17) and the more committed CD34+CD45ra+ progenitors (13.31% ± 8.16; Figure 3D) exhibited the lowest transfer rate. Instead, the multipotent progenitors (CD34+CD45ra-CD90+EPCR-, 20.02% ± 9.10) and stringently defined HSCs - CD34+CD45ra-CD90+EPCR+ were significantly more targeted than the parent populations (25.51% ± 11.19). These findings suggest a relationship between the phenotype and the frequency of mitochondrial transfer, which we further confirmed by identifying a positive correlation between the expression of the stem cell associated markers CD90+/EPCR+ and the mCherry mitochondrial mass transferred (**Figure 3E** and **F**).

**Figure 3.**
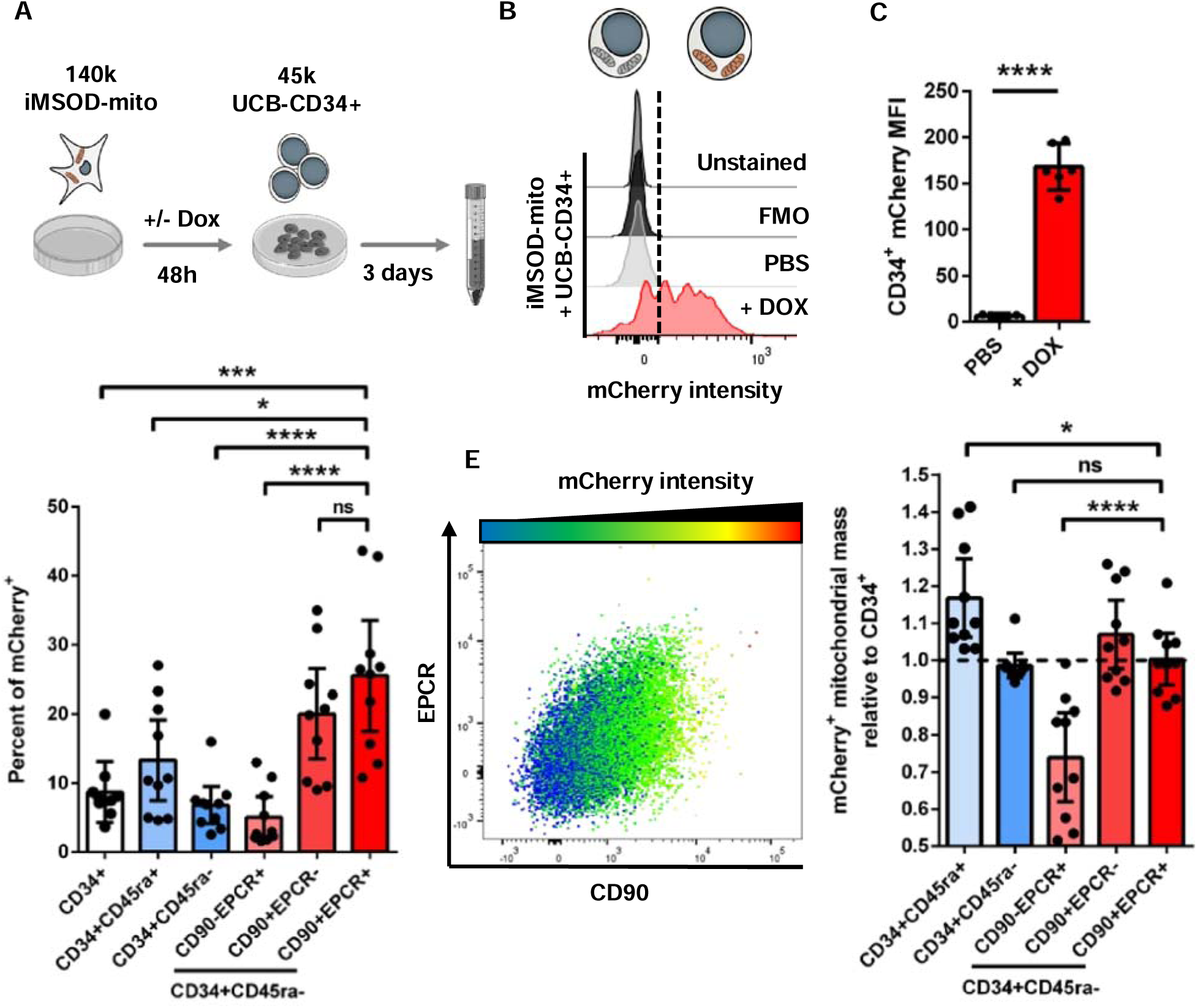
iMSOD-mito transfer mitochondria preferentially to phenotypically stem human HSPCs. **(A)** Experimental scheme of the coculture of human UCB-CD34+ cells on a confluent layer of iMSOD-mito with or without induction of mitochondrial mCherry tagging. **(B)** Flow cytometry analysis of mCherry intensity by flow cytometry reveal mitochondrial transfer from iMSOD-mito to human CD34+ after induction (+DOX). Controls consists of coculture with uninduced iMSOD-mito (PBS), unstained and fluorescence minus one (FMO)**. (C)** MFI of mCherry in UCB-CD34+ cells after coculture with (+DOX) or without (PBS) induction of iMSOD-mito. Unpaired t-test (n = 6). **(D)** Percentage of mCherry+ in hematopoietic subpopulations after coculture with induced iMSOD-mito. Unpaired t-test with logit transformation (n = 10). **(E)** Pseudo-color flow cytometry plot of mCherry intensity in CD34+CD45ra-according to EPCR and CD90 fluorescence intensity. **(F)** Transferred mCherry+ mitochondrial mass to hematopoietic subpopulations. Data are normalized to the transferred mCherry mitochondrial mass of CD34+ cells (dashed line). Unpaired t-test (n = 10). **** = *p*-value <.0001; *** = *p*-value <.001; ** = *p*-value <.01; * = *p*-value <.05; ns = *p*-value >.05.

In conclusion, we demonstrated here the mitochondria transfer of iMSOD-mito to healthy hematopoietic cells. Phenotypic HSCs were further identified as primary receiver of mitochondria, and mitochondrial mass correlates with expression of CD90+/EPCR+ markers.

### Primary human MSCs preferentially transfer mitochondria to HSCs in both 2D and 3D culture systems

To corroborate our findings with unmodified mesenchymal cells, primary BM-MSCs from three healthy donors were transduced with our mCherry-mitochondria labelling system (**Figure 4A, Supp. Figure 4A**), hereafter referred to as MSCs-mito. Using the same 2D co-culture experimental design, we first compared the transfer capacity of iMSOD-mito and MSCs-mito. Strikingly, while all primary MSC donors exhibited mitochondrial transfer capacity, this occurred at a reduced extent as compared to iMSOD-mito (3.38% ± 0.92 against 15.3% ± 2.25, **Figure 4B**). Interestingly, looking at the transfer in various hematopoietic populations revealed a similar biased clear increased frequency in HSCs. Despite an expected variability across donors, HSCs constantly received mitochondria the most among hematopoietic populations (5.87 to 19.1% of HSCs) in our 2D setting (**Figure 4C**, **Supp. Figure 4B**). As a comparison, CD34+ cells typically exhibited 2.76 to 4.92% of mCherry mitochondria (**Figure 4B**).

**Figure 4.**
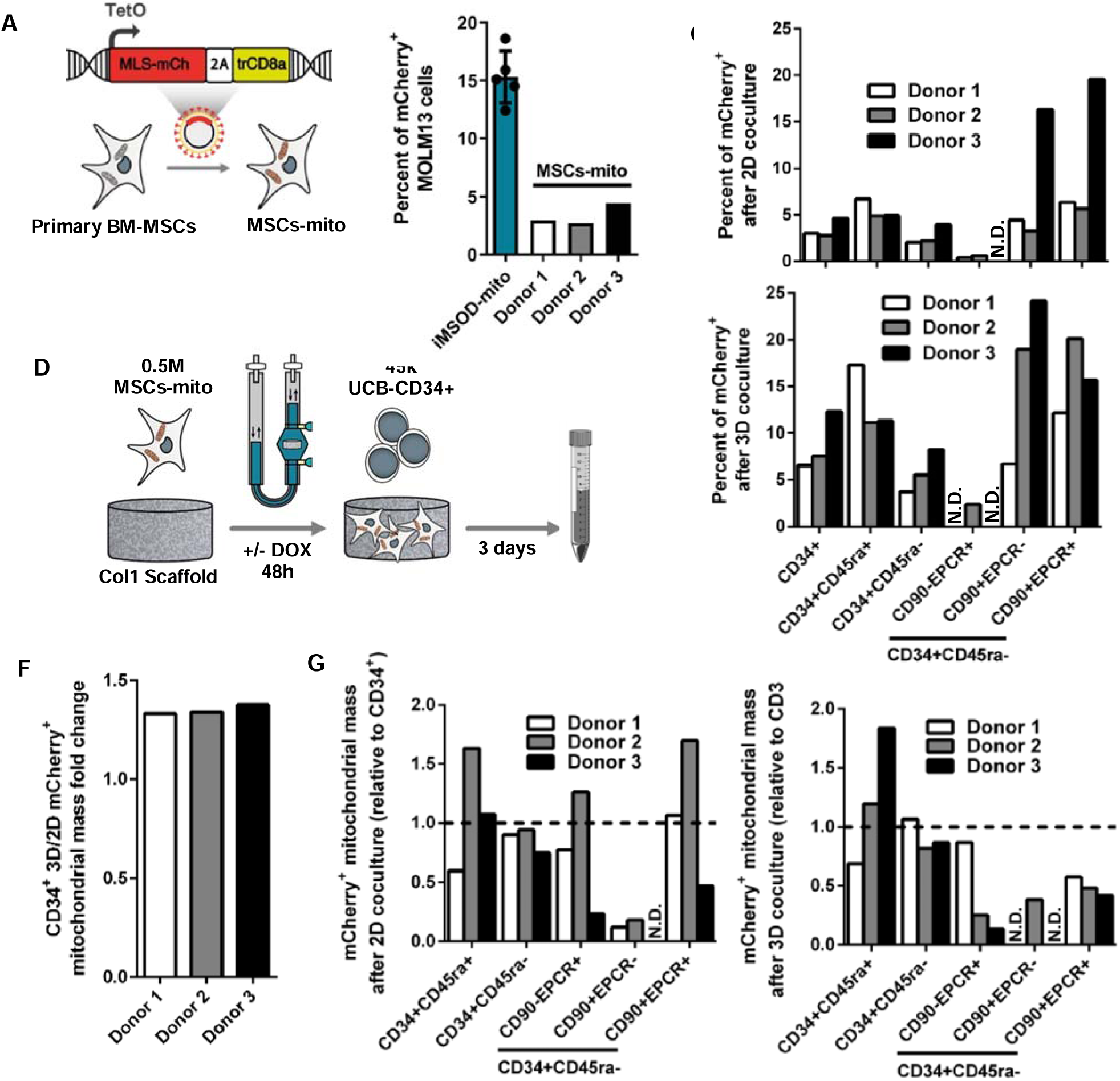
Primary human MSCs preferentially transfer mitochondria to HSCs in both 2D and 3D culture systems. **(A)** Diagram representing the genetic cassette transduced to the primary human bone marrow mesenchymal stromal cell (BM-MSCs) for the generation of the inducible mitochondria mCherry tagged MSCs-mito. **(B)** Percentage of mCherry+ MOLM13 cells after coculture with MSCs-mito or iMSOD-mito. **(C)** and **(E)** Percentage of mCherry+ in hematopoietic subpopulations after coculture with induced MSCs-mito in 2D and 3D respectively. **(D)** Experimental scheme of the 3D bioreactor coculture of UCB-CD34+ cells within a Collagen I scaffold colonized by MSCs-mito with or without induction of mitochondrial mCherry tagging. **(F)** 3D/2D fold change of transferred mCherry+ mitochondrial mass to CD34+ cells by MSCs-mito cells. **(G)** and **(H)** Transferred mCherry+ mitochondrial mass to hematopoietic subpopulations MSCs-mito cells in 2D and 3D respectively. Data are normalized to the transferred mCherry mitochondrial mass of CD34+ cells (dashed line).

We further enquire whether the establishment of mesenchymal niches in 3D perfusion bioreactor could impact the mitochondrial transfer (**Figure 4D**). After 3D co-culture with UCB-CD34+ cells, we observed a similar bias of transfer towards stem populations than in 2D. However, the 3D setting resulted in a greater percentage of mitochondria populations receiver, with a two-fold increase for HSCs (11.93 to 19.84% in 2D versus 3D respectively) and for CD34+ cells (6.57 to 12.3% in 2D versus 3D respectively, **Figure 4E**). Along with a higher percentage of transfer in 3D than in 2D, we observed a consistently greater mitochondrial mass transferred to CD34+ cells across all primary donors in 3D (1.35-fold increase ± 0.02; **Figure 4F**). However, the mitochondrial mass was rather heterogenous across hematopoietic subpopulations and no clear correlations could be established between 2D and 3D behaviors (**Figure 4G** and **4H**).

Of note, using iMSOD-mito for the engineering of 3D niches, we observed similarly an increased in the frequency of mitochondrial transfer in 3D in the CD34+ HSPCs (6.08% ± 1.82 in 2D versus 10.26% ± 0.74 in 3D, **Supp. Figure 4C**). This was also accompanied by a decrease in mitochondrial mass in the CD34+ receiving population (0.71-fold change ± 0.08. **Supp. Figure 4D**). Altogether, these findings establish clear similarities between the mitochondrial transfer occurring using primary MSCs and the iMSOD-mito, with a bias toward HSCs as primary receiver. Besides, 3D culture system was shown to significantly increase the mitochondrial transfer rate, which was previously not attenable by mesenchymal density increase in 2D.

Finally, we ambitioned to uncover potential functional consequences of the mitochondrial transfer. To this end, UBC CD34+ cells co-cultured with iMSOD-mito (in 2D or 3D) were sorted as mCherry+ and mCherry-populations and culture as single cells in 96 well plates for up to 7-day to assess division / quiescence activities (**Figure 5A**). A total of 192 cells were successfully isolated and followed at the single cell level in vitro. Viability assessment after two days first revealed a superior survival of CD34+ cells retrieved from 3D versus 2D conditions (**Supp. Figure 5A**). In 2D, CD34+ cells receiving mitochondria were more quiescent over the 7 days of culture compared to non-recipient CD34+ (**Figure 5B**). Indeed, after four days, 46.16% of cells remained undivided for CD34+mCherry+ in 2D, against 22.59% for mCherry-(**Figure 5C**). These findings were also observed for the second division: 60% of cells had one division for CD34+mCherry+ in 2D, against 52% for mCherry-at day 7 (**Supp. Figure 5B**). A similar trend was observed for cells retrieved from 3D conditions (**Figure 5B),** although the higher quiescence of CD34+mCherry+ was more marked for the second division (71% against 82% for CD34+mCherry-) (**Supp. Figure 5B**). Consistent with this, at the end of the 7-day clonal experiments, in 2D, a large fraction of mCherry+CD34+ remained undivided as compared to mCherry-CD34+ (26.67% versus 18.34% respectively) (**Figure 5C**) while no clear difference was observed in cells retrieved from 3D culture. Last, among dividing cells we seek to understand what conditions would lead to the largest colony formation (**Figure 5D**). In both 2D and 3D, mCherry+ cells were shown to possess higher colony formation capacity, especially within the 2D cultured cells with 80% remaining under 5 cells for mCherry+ cells against 96% for mCherry-(**Figure 5D**).

**Figure 5.**
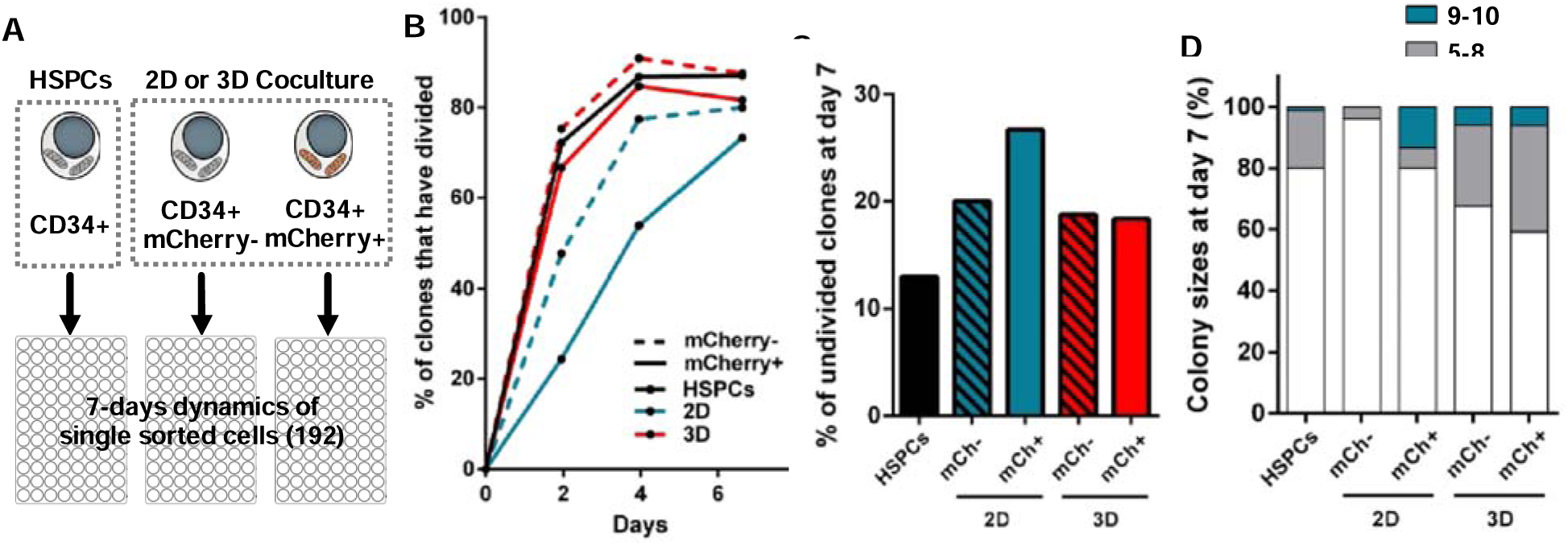
Mitochondria transfer is associated with increased quiescence in receiving blood populations. **(A)** Diagram representing the sorting of CD34+ after monoculture (HSPCs) and 2D and 3D coculture with iMSOD-mito for single cell clonal culture. **(B)** Evolution of the percentage of CD34+ clones which have undergone division overtime. **(C)** Percent of undivided CD34+ clones at day 7 of single cell culture. **(D)** Distribution of colony size from CD34+ clones at day 7 of single cell culture. N.D: not detected.

## Discussion

We here describe the iMSOD-mito as the first human mesenchymal line with inducible mitochondria labelling, exhibiting robust mitochondrial transfer to both malignant and healthy human hematopoietic cells. This transfer exclusively relies on direct cell-to-cell contact and is biased towards phenotypic HSCs. The engineering of 3D niches in perfusion bioreactor enhanced the mitochondrial transfer rate/mass resulting in apparent retained quiescence in receiving blood populations.

Beyond the standardization conferred by our human mesenchymal line, dye independent inducible mCherry-mitochondria labelling provides multiple advantages over conventional strategies to visualize mitochondrial transfer. First and foremost, as opposed to Mitotracker and equivalent chemical dyes, mCherry labelling is not cytotoxic and does not depend on the MMP (Neikirk *et al*., 2023). It can thus be exploited for longitudinal studies of the functional impact of the transfer as it does not disrupt mitochondrial biology. More importantly, mCherry intensity can be directly corelated to mitochondrial mass (de Almeida *et al*., 2017). It is also not excreted by the xenobiotic efflux pump which targets mitochondrial dyes and complicates their interpretation for mitochondrial transfer (de Almeida *et al*., 2017; Mansell *et al*., 2021). This is especially relevant for HSPCs, which are known to contain high levels of ABC xenobiotic efflux pumps (de Almeida *et al*., 2017) potentially leading to biased interpretation of transfer rates in stem populations. Lasty, the inducible expression of mCherry on the mitochondria allows for flexible experimental design, precise flow cytometry internal control, and low susceptibility to gene silencing.

We here evidenced high and robust mitochondrial transfer towards both leukemic and healthy HSPCs by iMSOD-mito. This standardized cell line was instrumental to uncover the preferential transfer towards HSCs and can be used to further discriminate the transfer mechanism leading to this preference. Genetic disruption could be performed to identify the complete molecular signaling at play before, during and after the mitochondria transfer, which current reports have yet to fully elucidate. Furthermore, as an unlimited human BM-MSCs cell source, iMSOD-mito could be used to screen new drugs which would target this mitochondrial transfer while preserving BM-MSCs integrity. This is of particular importance as mitochondrial transfer has been implicated in drug resistance mechanism in multiple leukemia models (Moschoi *et al*., 2016; Wang *et al*., 2018; Saito *et al*., 2021).

In sharp contrast to iMSOD-mito, primary BM-MSCs labelled with the same system exhibited lower and more variable mitochondrial transfer, a characteristic of primary BM-MSCs observed elsewhere (Polak *et al*., 2015). Primary BM-MSCs are known to display substantial phenotypic and functional differences across donors, but also within a sole individual due to high clonal diversity (Sacchetti *et al*., 2016). This higher transfer of iMSOD-mito compared to primary BM-MSCs can have multiple causes. First, iMSOD-mito is a clonal population, and thus the capacity to transfer mitochondria might be more uniformly distributed across all cells than its primary BM-MSCs “bulk” counterpart. Secondly, it was reported that BM-MSCs which adopt a cancer-associated fibroblast (CAF) phenotype provided more mitochondrial transfer to leukemic cells (Burt *et al*., 2019). The iMSOD-mito does not display morphological nor functional CAF characteristics, but the high telomerase expression ensuring immortalization is a common feature of cancer cells. Despite differences between iMSOD-mito and primary BM-MSCs, the same pattern of preferential transfer towards the phenotypically stem subpopulation of CD34+ HSPCs was observed across all donors, advocating for the iMSOD-mito capacity to mirror the transfer mechanism of primary cells. The conserved mechanism of transfer prompts the further exploitation of iMSOD-mito as a prime model to decipher this mechanism.

Mitochondrial transfer is typically related to stress conditions whereby the metabolic activity of recipient cells is rescued (Islam *et al*., 2012; Mistry *et al*., 2019). In the context of leukemia, leukemic stem cells can also sustain their high metabolic demands by hijacking mitochondria from BM-MSCs, thereby circumventing their own mitochondrial damages (Islam *et al*., 2012; Marlein *et al*., 2019; Mistry *et al*., 2019). However, among HSPCs, HSCs have the highest mitochondria mass per volume among all stem cells (de Almeida *et al*., 2017). This is counter-intuitive since HSCs in the bone marrow are pre-dominantly quiescent (Passegué *et al*., 2005; Takihara *et al*., 2019), thus with low metabolic needs. The reported accumulation of mitochondria in stem populations is in line with our in vitro study. We further demonstrate that transferred cells are more quiescent than non-recipient HSPCs. It is thus possible that our experimental setting reflects the in vivo mitochondrial assimilation by HSCs. This would give the opportunity to further study that process, and ideally identify the cellular signal from the receiving population as the most likely culprit for the transfer initiation. In fact, this remains largely elusive in the field, although a recent study reported a positive correlation between the level of CXC chemokine receptor 4 (CXCR4) expressed on recipient leukemic cells, and the rate of mitochondrial transfer from bone marrow MSCs (Giallongo *et al*., 2022). This is further correlated with the fact that HSCs have higher expression of CXCR4 than progenitors (Peled *et al*., 1999; Kollet *et al*., 2002; Triana *et al*., 2021)

While in 2D we show that one quarter of HSCs are transferred with mitochondria, the non-recipient pool contains 75% of the remaining phenotypic HSCs. While having a smaller pool of phenotypic HSCs we evidenced a retained quiescence for mCherry+ HSPCs. Moreover, this mitochondrial transfer within the niche could be a contributing mechanism to sustain such a high demand of functional mitochondria within HSCs.

We further propose the exploitation of 3D perfusion system to study the mesenchymal mitochondrial transfer. We previously established complex 3D bone marrow proxies using primary BM-MSCs, leading to robust maintenance of functional healthy HSCs (Bourgine *et al*., 2018) and expansion of primary leukemic material (García-García *et al*., 2021). Within our system, we observed a higher transfer rate in 3D as compared to 2D culture. Since mitochondrial transfer was shown to exclusively occur by short range TNTs establishment, we postulate that the 3D environment offered a higher surface of contact between cells and better recapitulate their native BM habitat. However, an increased iMSOD-mito density in 2D did not significantly impact the transfer efficacy. In 2D, the iMSOD-mito is grown as a monolayer and as such may not reflect the complexity and quantity of interactions occurring in 3D. Moreover, we previously reported a biased distribution of stem populations in the bioreactor chambers of 3D systems (Bourgine *et al*., 2018), while committed cells are released in the supernatant. Since mesenchymal cells are exclusively present in the chamber forming the niche, this could explain the higher mitochondrial transfer efficacy in our bioreactor device.

Taken together, the combined exploitation of iMSOD-mito and 3D culture systems bears high promises in multiple scopes. Fundamentally, this could facilitate high-throughput screening of molecules and thus identifying pathways modulating mitochondrial transfer efficacy. Translationally, such a platform could be harnessed to study mitochondrial disorders. Ideally, one can envision the revitalization of HSPCs by guided-mitochondria delivery, in line with the CD34+ mitochondrial augmentation of patients with single large-scale mitochondrial DNA deletion syndromes (Jacoby *et al*., 2022).

### Conclusion

In summary, our work lays the ground for the exploitation of iMSOD-mito for the standardized modeling of mesenchymal-to-hematopoietic mitochondrial transfer in 2D or 3D culture systems. This bears relevance to decipher the mechanism and function of such transfer in the human BM niche but may also apply to other organ systems whereby mesenchymal cells have key regulatory functions through mitochondrial delivery.

## Materials and methods

### Ethic statement

Experimental work carried out with primary human samples was approved by the regional and ethical committee for Lund/Malmö (Regionala Etikprövningsnämnden I Lund/Malmö), approval no. 2010-695. Informed consent was obtained from mothers of the umbilical cord blood (UBC) donors, and all samples were de-identified before use in the present study.

BM biopsies from the iliac crest bone were obtained from the Division of Pathology (Department of Clinical Sciences, Lund). Sampling and sample preparation were approved by the Institutional Review Committee in Lund (Regionala Etikprövningsnämnden I Lund, approval no. 2014/776 and 2018/86). All procedures and handling of human samples were performed in accordance with our approved ethical permits and the Helsinki Declaration of 1975, as revised in 2008.

### Bone marrow mesenchymal stromal cells expansion in 2D and 3D cell culture

2D culture of human BM-MSCs was carried out in a Nucleon™ Delta Surface 12-well plate (ThermoFisher #140675). Briefly, 144.000 MSOD cells were cultured for 2 days in complete media (CM) composed of 500 mL of MEM-α (Gibco # 22571038) supplemented with 10% of tetracycline-free fetal bovine serum (FBS; ThermoFisher); 5 mL of sodium pyruvate (100 mM; Gibco #11360070); 5 mL of HEPES (1 M; Gibco # 15630080); 5 mL of Penicillin-Streptomycin-Glutamine (100x at 50 mg/mL; Gibco #10378016) and ascorbic acid (AA; 100 µM; Sigma #A8960-5G). For mCherry mitochondrial labelling of the iMSOD-mito and MSCs-mito, 150 ng/mL of doxycycline hyclate (98%; ThermoFisher #15500554) was added to the CM. The murine BM-MSC line OP9 was cultured in CM with 20% FBS.

For 3D culture, cells were suspended at 0.5 million cells per 7 mL in CM with or without doxycycline. The bioreactor was mounted with a 6 mm diameter Collagen I scaffold (Avetene Ultrafoam, BD # 14648) as previously described (Dupard and Bourgine, 2021). After infusion of the 7 mL cells suspension, a first overnight infuse/withdraw perfusion cycle speed of 2.8 mL/min with displacement goal at 2 mL was performed to allow dynamic cell seeding to the scaffold; for the rest of the 3D culture the infuse/withdraw perfusion cycle speed was lower at 0.28 mL/min.

### Umbilical Cord Blood (UCB)-CD34+ cells isolation

UBC was collected at Skåne University Hospital and Helsingborg Hospital. Briefly, mononuclear cells were collected by Ficoll separation and CD34+ cells were isolated using the CD34 MicroBead kit (Miltenyi Biotec #130-046-702) according to the manufacturer’s instructions. UCB-CD34+ cells samples used in this study originated from a pool of a minimum of 3 units, processed within 24h after collection and with a minimum CD34+ purity of 94 % and viability of at least 95 %.

### Mesenchymal-Hematopoietic 2D and 3D coculture

The following was performed after BM-MSCs expansion for 2 days in addition of two washes with PBS.

For 2D coculture, carried out over 3 days, 45.000 UCB-CD34+ or 100.000 MOLM13 cells were added in each well in 1 mL of Coculture media (CoM), composed of DMEM (High Glucose, no glutamine, no calcium; ThermoFisher #21068028); 20% BIT9500 Serum substitute (StemCellTechnologies #09500), 1% Penicillin-Streptomycin-Glutamine (100x at 50 mg/mL, Gibco #10378016) and 1% HEPES (1 M; Gibco # 15630080). Media was supplemented of 0.02 % ß-mercaptoethanol (500X at 50 mM; ThermoFisher #31350010), human stem cell factor (SCF), thrombopoietin (TPO) and Fms-related tyrosine kinase 3 ligand (FLT3LG) at 10 ng/mL (all from Miltenyi Biotec, respectively #130-096-692, #130-095-745 and # 130-096-474) reconstituted in IMDM + 10% Bovine Serum Albumin (BSA; StemCellTechnology #09300). Media change was performed on day two.

For 3D, the same number of cells were resuspended in 5 mL and injected in the bioreactor as previously described (Dupard and Bourgine, 2021). After infusion of the 7 mL cells suspension, a first overnight infuse/withdraw perfusion cycle speed of 2.8 mL/min with displacement goal at 2 mL was performed to allow dynamic cell seeding to the scaffold; for the rest of the 3D coculture the infuse/withdraw perfusion cycle speed was lower at 0.28 mL/min.

### Sample preparation and flow cytometry

For 2D coculture, single-cell suspension was prepared by washing with PBS, followed by a 10 min digestion at 37 °C in Trypsin-EDTA (0.05%; Life Technology #25300054) supplemented with DNase I (0.25 mg/mL; Thermofisher #10700595). Digestion was stopped by the addition of an equal volume of CoM. For 3D culture, the liquid fraction was collected, and the bioreactor washed twice with PBS. The scaffold was incubated at 37°C in CM supplemented with 0.75 mg/mL of Collagenase (Sigma #C6885) for 15 min under agitation. Lysate was briefly trypsinated and pooled to the liquid fraction. Cells from 2D and 3D were then resuspended in FACS buffer composed of PBS with 2% FBS and 1 mM EDTA and passed through a 40 µm nylon mesh before staining. All samples were analyzed with an LSR Fortessa flow cytometer (BD Biosciences). Live-or-Dye™ 488/515Green (Biotum #32004) or DAPI (Sigma-Aldrich #D9542) were used according to the manufacturer’s protocol and allowed the exclusion of dead cells from the analysis. Monoclonal antibodies against human CD34 (BV421; 2:100; Clone: 581, BD #562577) and CD45ra (AF700; 10:100; BD #560673) allow the discrimination between committed (CD34+CD45ra+) and stem (CD34+CD45ra-) hematopoietic subpopulations. The stem cells compartment was further characterized with antibodies directed against EPCR (APC; 5:100; Biolegend #351905) and CD90 (PE-Cy7; 10:100; Clone: 5E10; Biolegend #561558). Phenotypic hematopoietic stem cells (HSCs) were identified with the gating CD34+CD45ra-CD90+EPCR+; Multipotent progenitor populations were defined as CD34+CD45ra- and further differentiated based on CD90 and EPCR expression (CD90+EPCR- and CD90-EPCR+). To identify MOLM13, we used the pan hematopoietic marker CD45 (AF700; 2:100; BD #560566). trCD8a expression on iMSOD-mito was measured with the monoclonal antibody CD8 (APC; Clone: RPA T8; BD # 555369). Samples were analyzed with FlowJo (version 10.7, BD Biosciences).

### Sorting and single cell clonal culture for quiescence assay

After monoculture or coculture, live HSPCs were sorted based on the expression of CD34 and their mCherry fluorescence intensity. The single cell sorting was performed with the FACSAria™ III Sorter (BD Biosciences) on 96 well plate (Sarstedt #83.3924.500) with high purity settings. The culture media consisted of CoM supplemented with SCF, TPO and FLT3 as described above with the addition of UM729 (1 µM; StemCellTechnologies #72332), 100 µL was used for culture. Visual cell counting and media change were performed every 2 days. On days 7 from cell sorting, the final counting was performed, and the experiment was stopped.

### Tracing of MOLM13 generations and mitochondrial transfer between iMSOD-mito

For tracing MOLM13 generations and iMSOD-mito to iMSOD-mito mitochondrial transfer we used Celltrace™ Violet (ThermoFisher # C34571) and Celltrace™ FarRed (ThermoFisher # C34572) cell proliferation kits respectively. Staining and acquisition was performed according to manufacturer’s protocol.

For MOLM13, stained cells were incubated for 3 days in coculture with doxycycline induced iMSOD-mito and generations determined by flow cytometry. For transfer of mitochondria between iMSOD-mito, two populations were incubated together. The first iMSOD-mito population was stained with CellTrace (CellTrace positive population) by was not induced with doxycycline. The second population was induced with doxycycline for two days but was not stained with CellTrace. After 24h of incubation of these two populations, mCherry was quantified in the CellTrace positive iMSOD population to determine the rate of mitochondrial transfer.

### Transduction of MSOD and primary MSCs-mito

Transduction of BM-MSCs cells was performed overnight via a third-generation lentiviral vector into which a doxycycline inducible polycistronic genetic cassette containing mCherry addressed to mitochondria with a mitochondria localization signal, and trCD8a separated with a 2A sequence. Viruses were produced by the Lund Stem Cell Center Vector Facility. For iMSOD-mito, clonal sorting based on expression mCherry and trCD8a was used to create a clonal cell line. To prevent exhaustion of the primary BM-MSCs, transduced cells were used without clonal sorting.

### Polymerase chain reaction (PCR) of murine and human mitochondrial and nuclear DNA

After cells lysis, nuclear DNA (nDNA) and mitochondrial DNA (mitDNA) from OP9 or UCB-CD34+ cells were collected with the Quick-gDNA™ Miniprep Kit from Zymo Research. DNA was then amplified with human or mouse specific primers for nDNA and mitDNA. For human mitDNA, primers used were FWD 5’ CCTTCTTACGAGCCAAAA 3’ and REV 5’ CTGGTTGACCATTGTTTG 3’; while for mouse, the primer used were: FWD 5’ ATCTGCTTCAATAATTTAATTTCACT 3’ and REV 5’ TTGTGAGTAGAAGTAAAATAATAAATGTAATGG 3’. For nDNA, human FWD primer was 5’ GCTGCTTCTCATTGTCTCGG 3’, and mouse FWD primer was 5’ CCTGCTGCTTATCGTGGCTG 3’. For both mouse and human nDNA, the REV primer was 5’ GCCAGGAGAATGAGGTGGTC 3’. The PCR products were run on a 1.5% agarose gel at 100V for 1 hour. The gel was subsequently acquired with BioRad Gel Doc XR+ Imaging System and analyzed using ImageJ.

### Immunofluorescence staining and live imaging

For visualization of the mCherry expression, iMSOD cells were grown on a coverslip for 24h, fixed for 20 min in 4% paraformaldehyde and counterstained with a DAPI solution at 0.1 µg/mL. EGFP, mCherry and DAPI were then acquired with the Leica Stellaris 5 Confocal Laser-Scanning Inverted Microscope. For colocalization of mCherry and TOMM20, we used a rabbit anti-TOMM20 monoclonal antibody (Abcam #ab56783) and a rat anti-tubulin antibody (Abcam #ab6160) for confirming cellular integrity after fixation. For density assessment of the iMSOD-mito culture, EGFP from live cells was acquired and manual counting was performed on three fields of view. For live imaging, 10.000 iMSOD-mito cells with or without 2 days of prior doxycycline induction were seeded onto 8-well glass bottom plate (ibidi #80827) in 300 µL of CoM without phenol (DMEM; Gibco #21063029). iMSOD-mito were left at 37°C overnight and 10.000 MOLM13 cells were added immediately prior to acquisition with a Nikon Ti2 widefield microscope. Acquisition of EGFP, mCherry and DIC was performed every 5 min for 12h to visualize mitochondrial transfer.

### MMP and ROS activity measurements

Mitochondrial membrane potential (MMP) was analyzed using the mitochondrial dye DiOC6(3). Prior to flow cytometry acquisition, MOLM13 cells from monoculture or coculture were incubated for 30 min with 20 nM of DiOC6(3) and 50 nM of Verapamil (Sigma-Aldrich #V4629) in phenol-free DMEM. Controls for mitochondrial staining specificity were acquired after 30 min incubation with the mitochondria uncoupler FCCP (300 nM; Sigma-Aldrich #C2920).

For reactive oxygen species analysis, similarly as for MMP, cells were analyzed after 30 min incubation with 50 µM of DCFDA / H2DCFDA (Cellular ROS Assay Kit; Abcam #ab113851). Staining specificity was confirmed with cell exposure to tert-Butyl hydroperoxide solution according to manufacturer’s protocol.

### Statistical analysis

Statistical analysis was performed with R (version 4.1.3) and GraphPad Prism (Version 9). Tests used are described in figure legends throughout the manuscript, with prior validation of test assumptions.

## Author Contributions

S.J.D. conceptualized the study. S.J.D., J.M.P. and P.E.B. designed the experiments and interpreted the results. S.J.D. contributed to all experiments described in the manuscript. J.M.P. performed and assisted the cocultured experiments, the colocalization confocal acquisition and analysis. A.G.G. and S.K. provided support for coculture experiments. A.B., D.Z. and D.H.G. isolated and expanded the primary human BM-MSCs. A.B. assisted the sorting of CD34+ cells. V.S. performed the setup and the acquisition of the live imaging mitochondrial transfer. S.J.D. wrote the initial draft. S.J.D. and P.E.B. wrote the final manuscript. All authors contributed to the manuscript editing.

## Acknowledgments

We thank Lund Stem Cell Center FACS Facility, Imaging Facility and the Lund Stem Cell Center Vector Facility. Lund University is gratefully acknowledged for experimental resources. S.J.D. would like to give special thanks to Jonas Larsson, Darcy Wagner, Niels-Bjarne Woods, Anna Fossum, Emanuela Monni, and Ani Grigoryan for their support and advice towards the completion of this study.

---

**Supplementary Figure 1.**
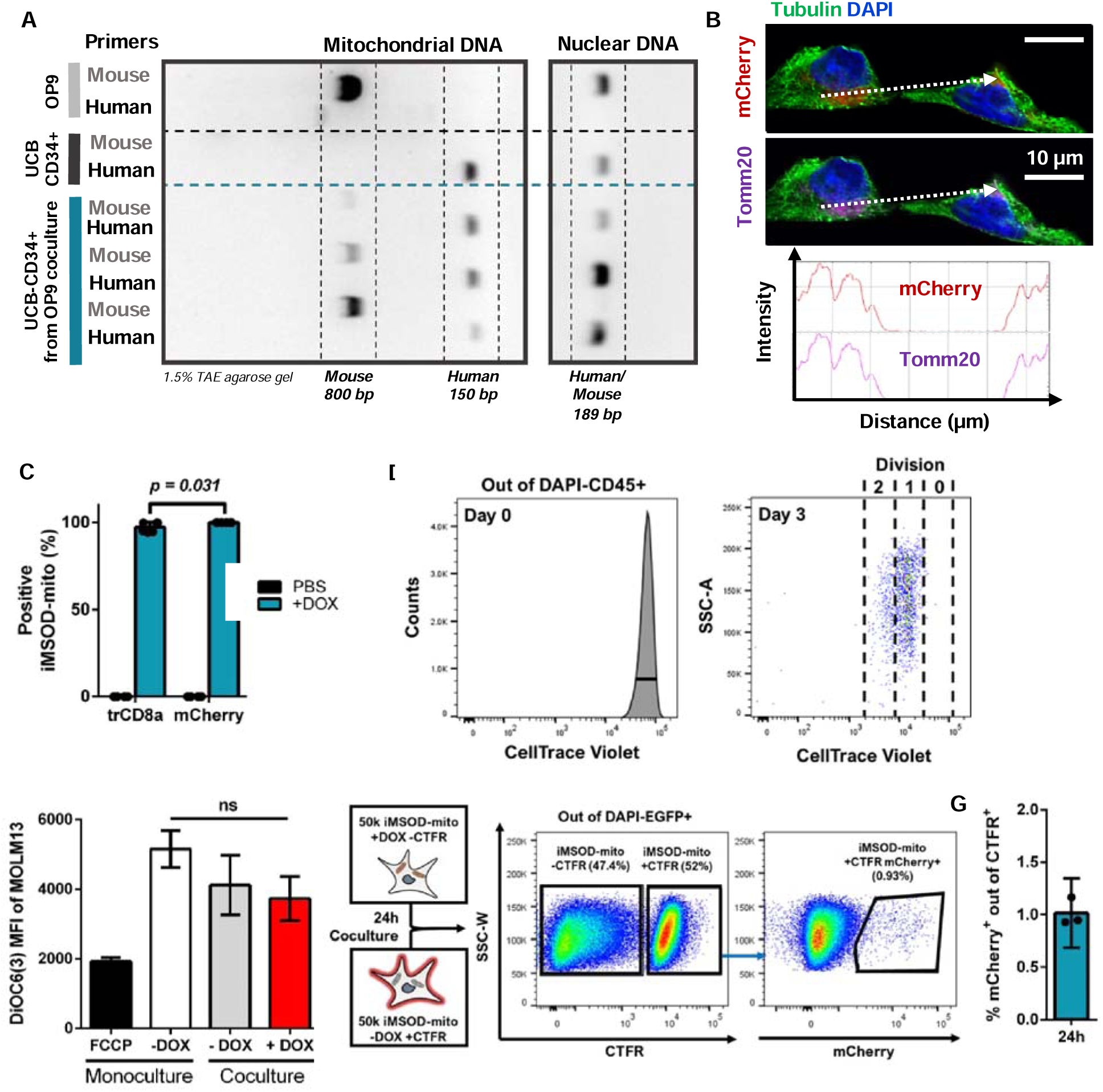
(A) The mouse mesenchymal stromal cell line OP9 and human umbilical cord blood (UCB) CD34+ cells were coculture for 3 days. CD34+ were sorted by flow cytometry and their DNA was extracted. Polymerase chain reactions (PCRs) with pairs of primers specific for either human or mice mitochondria DNA and nuclear DNA were used to identify mitochondrial transfer. Controls consist of DNA extracted from OP9 or UCB-CD34+ in monoculture. PCR products were visualized by agarose gel electrophoresis. (B) Confocal microscopy of iMSOD-mito after 24h exposure to 150 ng/mL of doxycycline staining with TOMM20. mCherry and TOMM20 fluorescence intensity along the same cross-section (dashed arrow) was used to determine fluorescence colocalization (C) Percentage of iMSOD-mito positive for mCherry and trCD8a after induction (+DOX) or no induction (PBS). Unpaired t-test after logit transformation (n = 4). (D) Flow cytometry gating of MOLM13’s divisional history according to the intensity of CellTrace Violet after 3 days of coculture with iMSOD-mito. (E) Mitochondrial membrane potential of MOLM13 cells measured by DiOC6(3) mean fluorescence intensity after monoculture or coculture with iMSOD-mito. Carbonyl cyanide-p-trifluoromethoxyphenylhydrazone (FCCP) mitochondrial depolarization confirms mitochondrial specificity of DiOC6(3) staining. One-way ANOVA (n=3). ns = p-value >.05. (F) Mitochondrial transfer analysis between uninduced (-DOX) iMSOD-mito stained with CellTrace Far Red (CTFR+) were placed in coculture with induced (+DOX) but not stained (CTFR-) iMSOD-mito for 24h. Mitochondrial transfer was identified when mCherry was detected in CTFR+ iMSOD-mito cells. (G) Percentage of mCherry+CTFR+ iMSOD-mito cells after coculture. (n = 3).

**Supplementary Figure 2.**
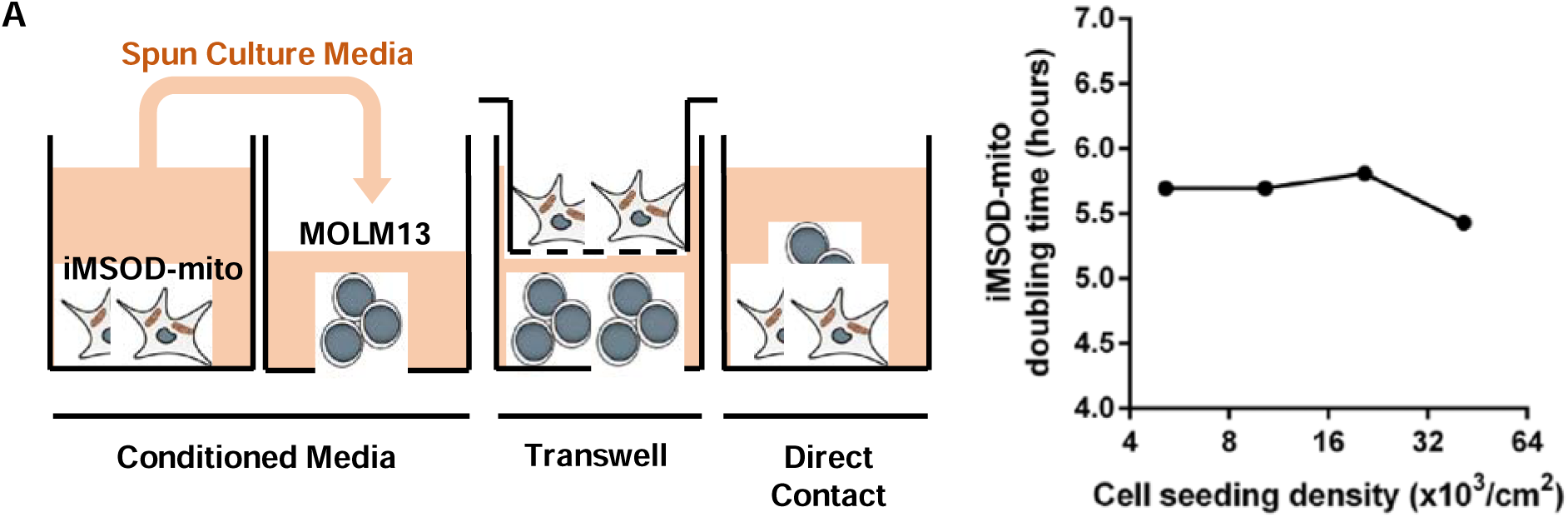
(A) Diagram of the different coculture strategies used to determine the mode of transfer of mitochondrial from iMSOD-mito to MOLM13 cells. The conditioned media approach consists of exposing MOLM13 to the supernatant collected from the media of a two-day induction of iMSOD-mito, spun at 400g for 5 min. In transwell, iMSOD-mito and MOLM13 were physically separated by a porous membrane. (B) Doubling time of iMSOD-mito over a culture of 3 days starting at different seeding densities.

**Supplementary Figure 3.**
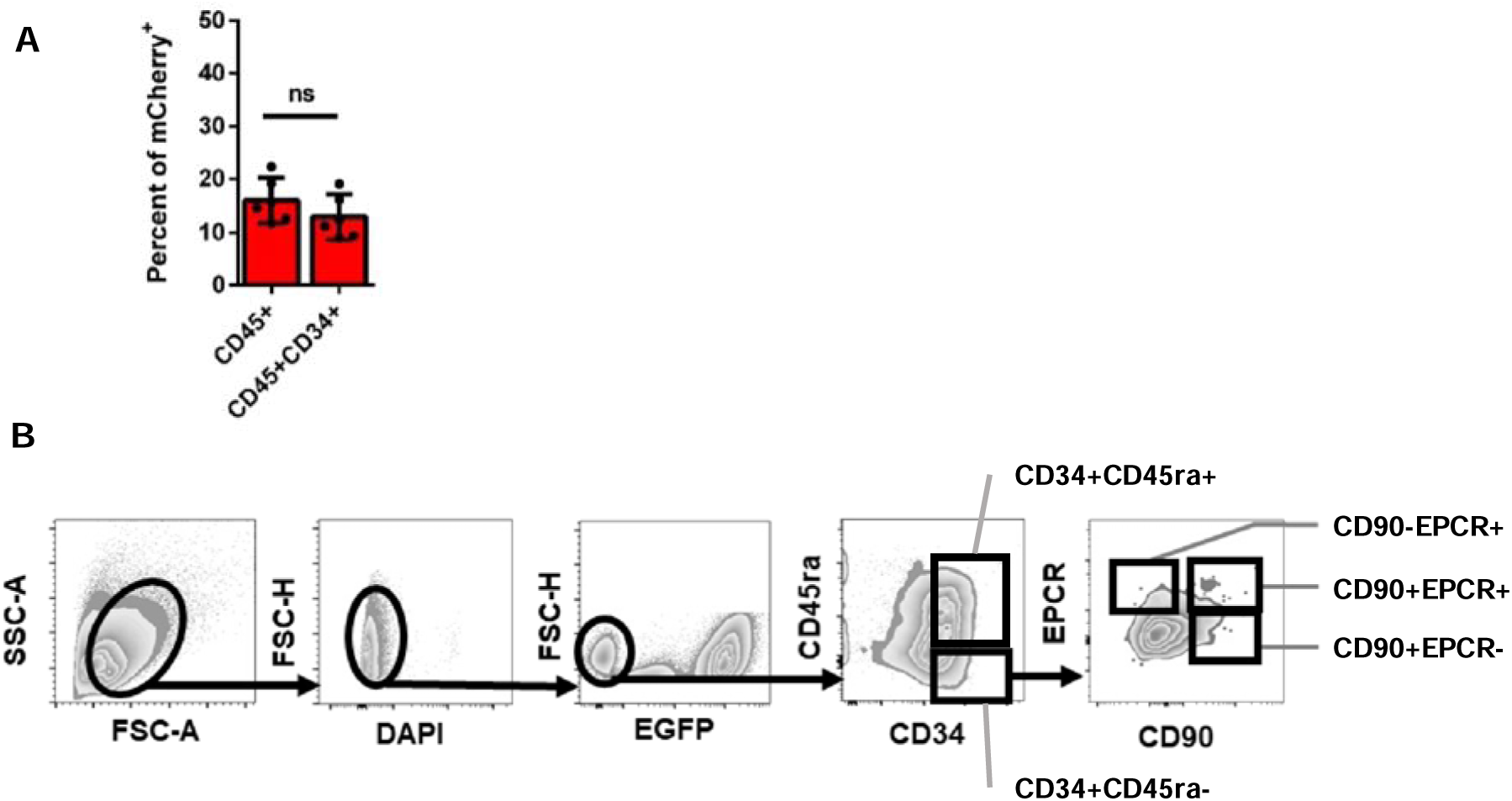
(A) Percentage of mCherry+ within the pan hematopoietic marker CD45+ cells and the pan stem and progenitor CD34+ cells after coculture with induced iMSOD-mito. Unpaired t-test with logit transformation (n = 4). ns = *p*-value >.05. (B) Diagram of the flow cytometry strategy to isolate hematopoietic stem and progenitor cells subpopulations.

**Supplementary Figure 4.**
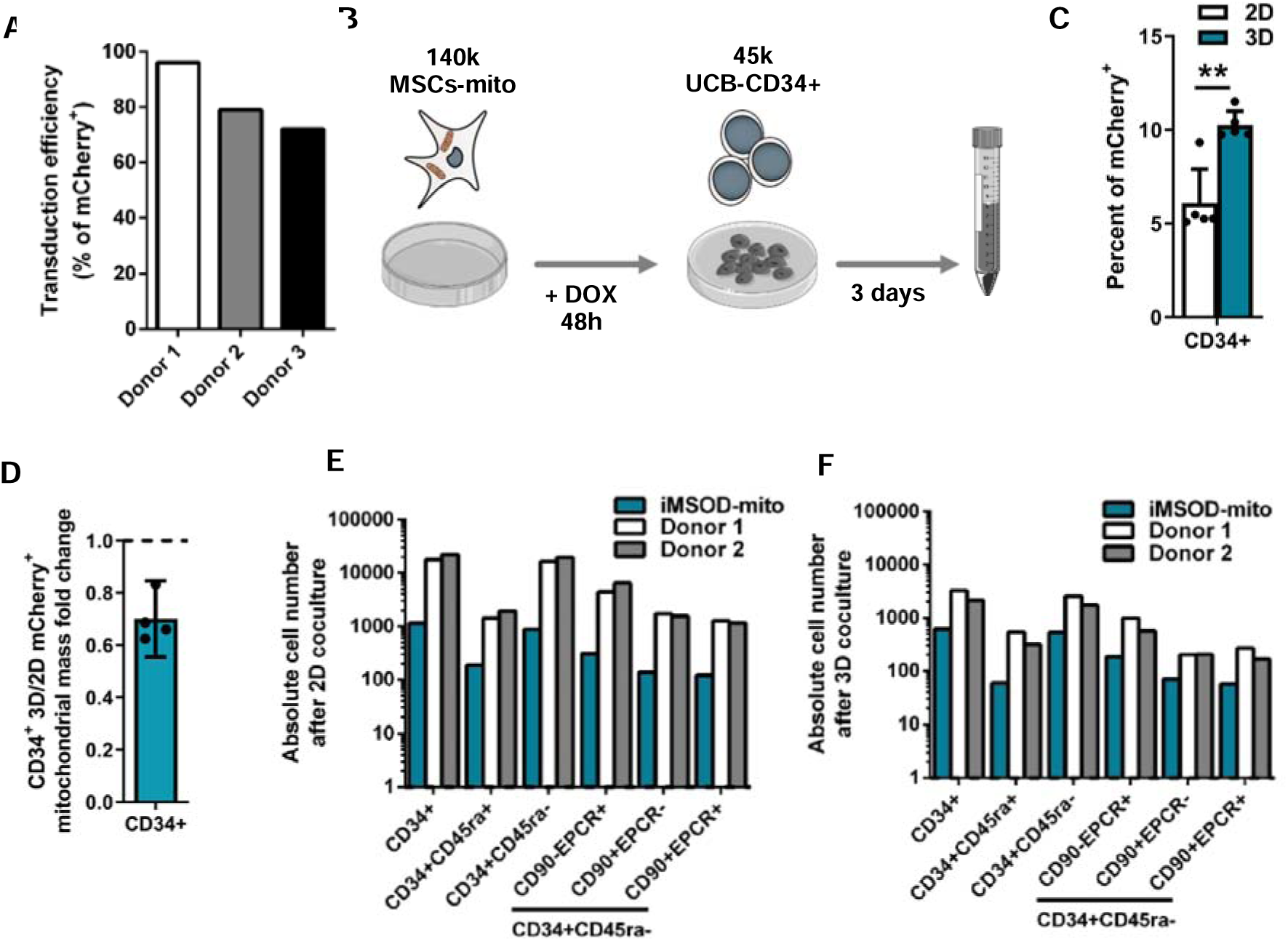
(A) Percent of mCherry+ MSCs-mito among donor after transduction and induction with 150 ng/mL of doxycycline. (B) Experimental scheme of the coculture of human UCB-CD34+ cells on a confluent layer of MSCs-mito with or without induction of mitochondrial mCherry tagging. (C) Percentage of mCherry+ in CD34+ after coculture with induced iMSOD-mito in 2D and 3D. Unpaired t-test with logit transformation (n = 5). ** = *p*-value <.01. (D) 3D/2D fold change of transferred mCherry+ mitochondrial mass to CD34+ cells by iMSOD-mito cells. (E) and (F) Absolute number of cells recovered across hematopoietic subpopulations after coculture with MSCs-mito or iMSOD-mito in 2d and 3D respectively.

**Supplementary Figure 5.**
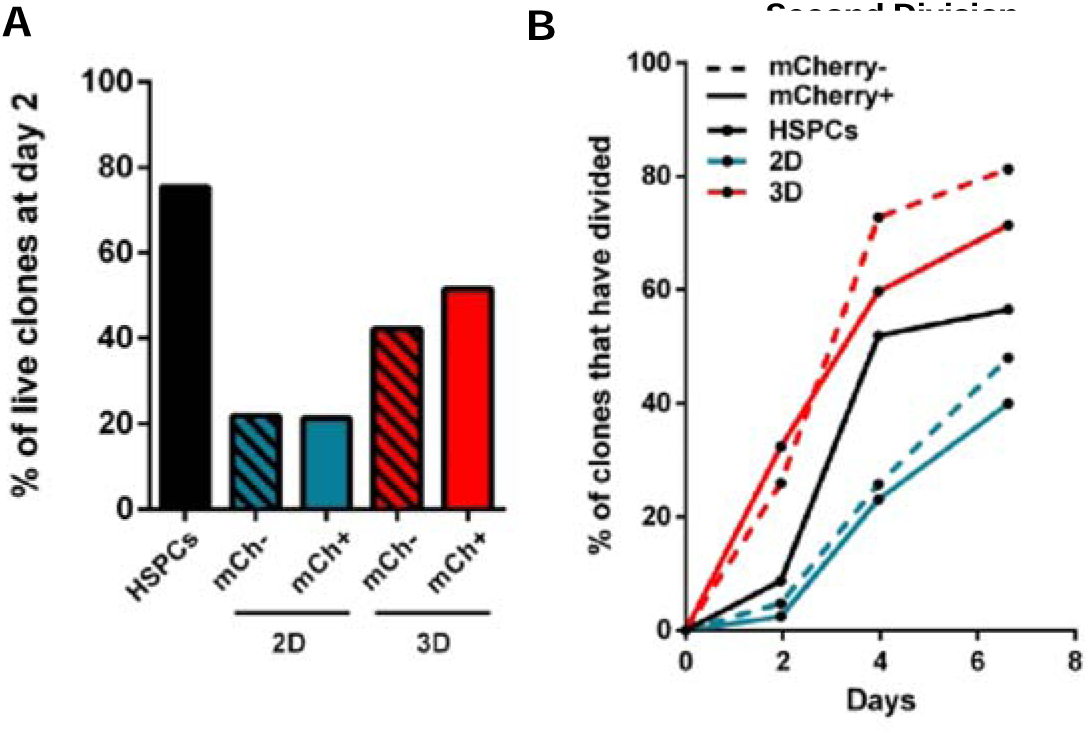
(A) Percentage of survival among sorted CD34+ clones at day 2. (B) Evolution of the percentage of CD34+ clones which have undergone their second division overtime.

